# When expectations are not met: unraveling the computational mechanisms underlying the effect of expectation on perceptual thresholds

**DOI:** 10.1101/545244

**Authors:** Buse M. Urgen, Huseyin Boyaci

**Affiliations:** Interdisciplinary Neuroscience Program, Bilkent University, Ankara, Turkey; Aysel Sabuncu Brain Research Center & National Magnetic Resonance Research Center, Bilkent University, Ankara, Turkey; Department of Psychology, Bilkent University, Ankara, Turkey; Department of Psychology, Justus Liebig University Giessen, Giessen, Germany

**Keywords:** expectation, visual perception, perceptual inference, Bayesian model, predictive computation

## Abstract

Expectations and prior knowledge strongly affect and even shape our visual perception. Specifically, valid expectations speed up perceptual decisions, and determine what we see in a noisy stimulus. Bayesian models have been remarkably successful to capture the behavioral effects of expectation. On the other hand several more mechanistic neural models have also been put forward, which will be referred as “predictive computation models” here. Both Bayesian and predictive computation models treat perception as a probabilistic inference process, and combine prior information and sensory input. Despite the well-established effects of expectation on recognition or decision-making, its effects on low-level visual processing, and the computational mechanisms underlying those effects remain elusive. Here we investigate how expectations affect early visual processing at the threshold level. Specifically, we measured temporal thresholds (shortest duration of presentation to achieve a certain success level) for detecting the spatial location of an intact image, which could be either a house or a face image. Task-irrelevant cues provided prior information, thus forming an expectation, about the category of the upcoming intact image. The validity of the cue was set to 100, 75 and 50% in different experimental sessions. In a separate session the cue was neutral and provided no information about the category of the upcoming intact image. Our behavioral results showed that valid expectations do not reduce temporal thresholds, rather violation of expectation increases the thresholds specifically when the expectation validity is high. Next, we implemented a recursive Bayesian model, in which the prior is first set using the validity of the specific experimental condition, but in subsequent iterations it is updated using the posterior of the previous iteration. Simulations using the model showed that the observed increase of the temporal thresholds in the unexpected trials is not due to a change in the internal parameters of the system (e.g. decision threshold or internal uncertainty). Rather, further processing is required for a successful detection when the expectation and actual input disagree. These results reveal some surprising behavioral effects of expectation at the threshold level, and show that a simple parsimonious computational model can successfully predict those effects.

## Introduction

Conventional models of perception postulate that perception is a process which is implemented by the bottom-up processing in the brain, where the physical properties of a stimulus is processed by different levels of the cortical hierarchy with increasing complexity. However, in a dynamic, contextually rich environment with an often ambiguous input, the visual system cannot process all sensory information accurately at once in detail. To decrease the computational burden of this process higher level mechanisms have been suggested to be involved in the information processing, which make our decisions become faster and more efficient (Summerfield and Egner, 2009; Summerfield and De Lange, 2014). For instance, while we are searching for a painting in a room, we look at the locations where the painting is more likely to be placed, i.e. the wall, instead of searching every single item/place in the room. Or when the sensory information we experience is ambiguous or noisy, it may be sometimes difficult to recognize the stimulus because there may be several interpretations of it. However, we usually come up with a single interpretation very quickly, because our prior knowledge facilitates perception while making decisions (Bar, 2004; Kok et al., 2012; Summerfield and De Lange, 2014).

Accordingly, computational models that posit perception as an inference process emphasize the role of top-down effects of prior information on perceptual decisions (Rao and Ballard, 1999; Friston, 2005; Heeger, 2017). Consistent with these models, empirical findings have confirmed that perception is not a process solely determined by the bottom-up processing in the brain. While the low-level properties of a stimulus is processed, there is also a top-down influence (i.e. context-based) on the perceptual processing from higher levels of the cortical hierarchy (Bar, 2004; Gilbert and Sigman, 2007; Summerfield and Koechlin, 2008; Muckli and Petro, 2013; Muckli et al., 2015; de Lange et al., 2018). It is by now well-established that visual perception is a process which results from an interplay of bottom-up and top-down information processing.

Bayesian models of perception provide a mathematical framework for this inference process (Teufel et al., 2013). Specifically, because the sensory information we experience is often ambiguous or noisy, the system combines the incoming sensory input with the prior to decide on the most probable causes of the sensory input (Mamassian et al., 2002; Kersten et al., 2004; Yuille and Kersten, 2006; Maloney and Mamassian, 2009; Summerfield and De Lange, 2014; de Lange et al., 2018). This is why the system can reliably make a decision although there are several interpretations of a sensory input or very similar retinal input may result in totally different percepts. Several work support the idea that perception is a probabilistic inference process, where the perceptual decisions are made by combining the priors with the statistical regularities in the environment (Weiss et al., 2002; Kersten et al., 2004; Knill and Pouget, 2004; Yuille and Kersten, 2006; Chalk et al., 2010; de Lange et al., 2018). This indicates that under some circumstances the system makes optimal interpretations and human behavior may approximate Bayesian ideal observer (Ernst and Banks, 2002; Kersten et al., 2004; Yuille and Kersten, 2006).

Accordingly, a growing body of literature have revealed that *expectations* that are formed based on our prior knowledge can bias perceptual decisions (Weiss et al., 2002; Sterzer et al., 2008; Summerfield and Koechlin, 2008; Summerfield and Egner, 2009; Chalk et al., 2010; Kok et al., 2011; Sotiropoulos et al., 2011; Kok et al., 2012; Wyart et al., 2012; Kok et al., 2013; Summerfield and De Lange, 2014; de Lange et al., 2018). Empirical findings which reveal the role of expectations on perception mainly come from the perceptual decision-making studies where reaction time is commonly used as the measure, which is an index of both perceptual and decision-making processes. It is found that expected stimulus (or in a cued-paradigm *congruent* stimulus) is detected *faster* and *more accurately* than the unexpected (*incongruent*) stimulus (Wyart et al., 2012; Stein and Peelen, 2015). Even though the role of expectations on perceptual decisions has gathered considerable support from these studies, the computational mechanisms giving rise to such a difference in detecting or recognizing the expected and unexpected stimuli remain unclear. In this study, by measuring perception at threshold level we aim to investigate how expectations affect *early visual processes*, which is distinct from motor and cognitive components of a decision-making process. We specifically investigate whether expectation has an effect on detecting the spatial location of a stimulus (also called *individuation*) while systematically manipulating the expectation validity in different experimental conditions. We measure duration thresholds, which is the shortest duration of the presentation that participants can successfully determine the location of the stimulus. Next, we present a *recursive* Bayesian updating scheme in which the prior is not fixed, but updated at each iteration to model the empirical results of the current study. Our findings expand on the behavioral effects of expectation on low level visual processing by unraveling the computational mechanisms that underlie the perceptual effects we found. We also discuss our findings within the framework of predictive computational models.

## Behavioral Experiment

### METHODS

#### Participants

Eight naive participants (4 female; 24.5 *±* 2.33 years) participated in the behavioral experiment that included four separate experimental conditions. All participants had normal or corrected to normal vision and reported no history of neurological disorder. Participants gave their written informed consent prior to the experiment. The experiment was approved by the Research Ethics Committee at Bilkent University.

#### Stimuli

Stimuli consisted of two category of images: ten face images (five female; age range was 19-69) taken from Face Database of the Park Aging Mind Laboratory (Minear and Park, 2004) and ten house images from Scene Understanding Database from the Princeton Vision Group (Xiao et al., 2010). Cues (*informative*: face, house; *uninformative* (neutral): question mark) used in different experimental conditions were taken from The Noun Project’s website (www.thenounproject.com; House by OCHA Visual Information Unit, Person by Alex Fuller, Question by Vicons Design from the Noun Project) and were scaled to 3.5 x 3.5*°* visual angle. As mask, scrambled version of the images were generated by dividing the image into 49 cells via creating 7 x 7 grids for each. After that each cell was randomly assigned to different locations. The stimuli including intact images (target stimuli) and mask images were scaled to 10.5 x 10.5*°* visual angle, converted to grayscale, histogram-matched (scaled to the mean luminance of all stimuli) by using SHINE Toolbox (Willenbockel et al., 2010), and adjusted and matched to a very low contrast value (2%). Experiments were programmed in MATLAB 2016a using Psychtoolbox (Brainard, 1997). Stimuli were shown on a CRT monitor (HP P1230, 22 inches, 1024 x 768 resolution, refresh rate 120 Hz.)

#### Experimental Design

Stimuli were presented on a gray background (RGB: 128, 128, 128). Each trial started with a cue simultaneously with a fixation dot located on the center of the cue, and presented for 2 seconds at the center of the screen. Cues were either *informative* (face and house) or *neutral* (question mark) depending on the experimental condition (See *Experimental Session* for details). Next, a target stimulus, which was an intact face or house image, and a scrambled version of the same image were simultaneously shown in left and right side of the cue at 10*°* eccentricity. Presentation duration of these images were determined by an adaptive staircase procedure (See *Procedure* for details). Next, as masks, different scrambled versions of that target stimulus were shown on the same locations for 64 ms. Following this, an empty display with a gray background was presented until a response is given by the participants. Participants’ task was to detect the spatial location of the target stimulus as soon as possible by pressing the left or right arrow key of the keyboard while maintaining their fixation on the fixation dot during the trial. Finally, a feedback message was given as “correct” or “wrong” to the participants for 752 ms. When the category of the cue and the image is the same, these trials are called *congruent* (expected) trials. When the category of the cue and the image is different, these trials are called *incongruent* (unexpected) trials. Note that equal number of each cue (face and house) appeared in random order in the experimental conditions where an informative cue is presented. Also note that equal number of each target stimulus (face and house image) was presented in all experimental conditions, and the target stimulus was randomly assigned to one of the two locations (left or right) in each trial. See Figure 1 for sample trials from the experiment.

**Figure 1:**
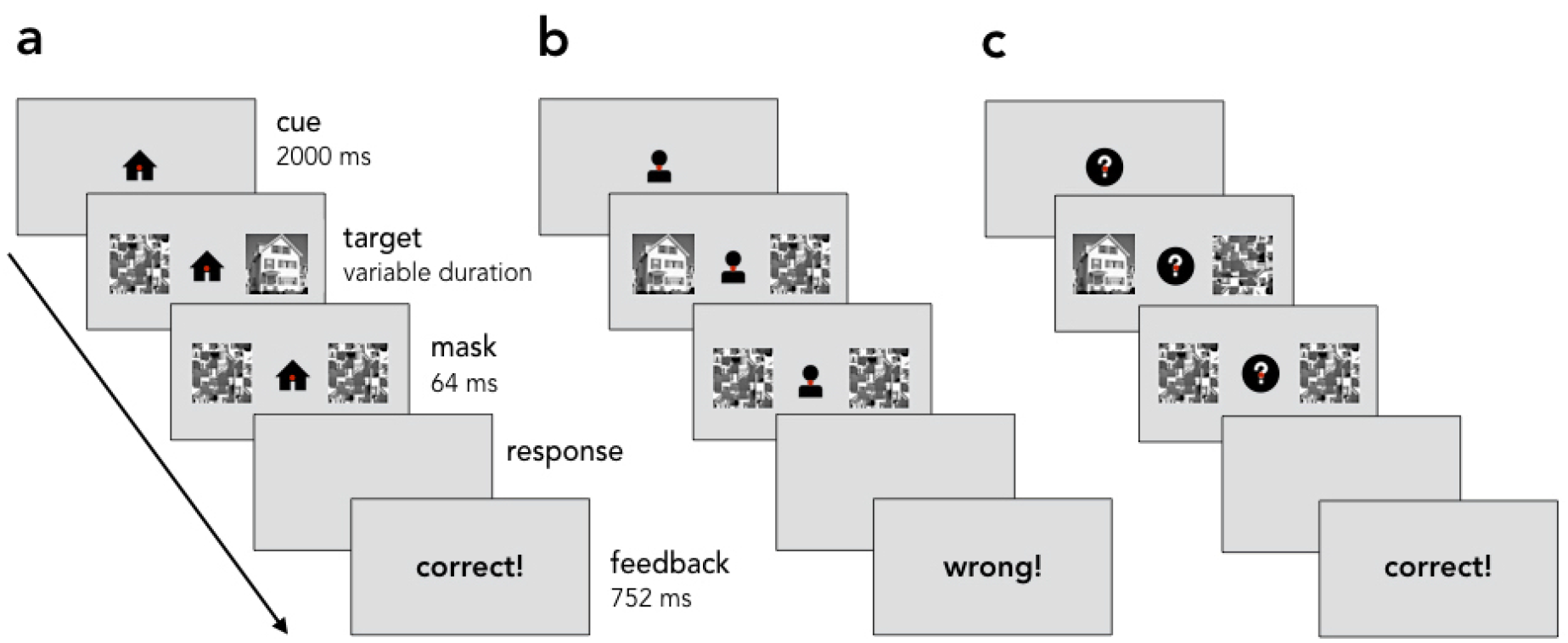
Behavioral experiment. Sample trial sequences. a. Congruent trial. b. Incongruent trial. c: Neutral trial. In all but the neutral trials a centrally presented cue predicted the category of upcoming target with a certain validity (100, 75, and 50%). Duration threshold, which is the minimum duration required to successfully detect the location of the target intact image, is determined per participant under each condition. See text for more details.

#### Procedure

Behavioral experiment consisted of a training session and an experimental session which comprises four experimental conditions^1^. In both sessions, 2-down 1-up adaptive staircase procedure with a two alternative forced-choice (2-AFC) paradigm was applied to derive duration thresholds (70.7% accuracy) in different trial types: *neutral* trials, *congruent* trials and *incongruent* trials in different conditions (see *Experimental Session* for details). Presentation duration of the target image and scrambled version of it were varied adaptively from trial to trial. The duration of each trial was determined by the accuracy of the participants’ responses in previous trials. Specifically, each wrong answer or two consecutive correct answers resulted in approximately 8 ms (step size) increase or decrease of the duration of the following trial target presentation respectively. At the beginning of each experimental condition, one staircase started from a relatively short duration (varied for each participant, minimum 8 ms), and the other staircase started from a relatively long duration (varied for each participant). There were 30 trials in each staircase in all experimental conditions, but number of staircases varied for each experimental condition.

#### Training Session

Prior to the experimental session, each participant completed a training session in order to stabilize their perceptual thresholds. Participants were seated 60 cm away from the screen and their heads were stabilized with a chin-rest. The training session consisted of 2 to 5 short experiments where the cue was always informative (face and house cue) and 100% valid in indicating the target stimulus category. Each experiment in the training phase had 120 trials and there were equal number of face and house cue trials. Number of experiments completed in the training phase varied for each participant, and it is determined by whether the participant’s threshold stayed within an interval of 8 ms (step size) for at least two sequential experiments.

#### Experimental Session

All participants completed four experimental conditions in randomized order in separate sessions. Participants were informed about the cue-validity prior to the experiments. Cue validity refers to the probability that the cue correctly predicts the category of the upcoming intact image.

### 100%-validity condition

In this experimental condition the cue (face or house) informed participants about the upcoming target stimulus category (either face or house image) with a 100% validity so that there was no violation of expectations. There were 120 (congruent) trials in total including 60 trials where the target was a face image following a face-cue, and 60 trials where the target was a house image following the house-cue.

### 75%-validity condition

In this experimental condition the cue informed about the correct category of the intact image with 75% probability (face or house). Equal number of each cue (face and house) were presented, and there were 480 trials in total. There were 360 *congruent trials* where the image category was correctly predicted by the cue, and 120 *incongruent trials* where the cue misled the participants about the upcoming image category.

### 50%-validity condition

In this experimental condition the cue validity was at 50%. Therefore, in total there were 240 trials, of which 120 were *congruent* and 120 were *incongruent*. Equal number of each cue was presented.

### Neutral (no expectation) condition

This experimental condition was included as a control condition because there was no informative cue (face or house) that informs participants about the upcoming image category. Rather, the cue was *neutral*, a question mark, during the experiment. Therefore, expectations about the upcoming stimuli were not formed. Except the cue type, all experimental stimuli and design were the same as the other conditions. There were 120 trials in total, and equal number of each image category was presented.

#### Statistical Analysis

Duration thresholds (70.7% accuracy) for spatial location detection in congruent, incongruent and neutral trials were estimated using the Palamedes toolbox (Kingdom and Prins, 2010) with Logistic function using Matlab 2016a. A 2 (congruency: congruent, incongruent) x 2 (validity: 75%, 50%) repeated measures ANOVA was conducted to investigate the effect of expectation on duration thresholds. Also, we conducted two-sample paired *t*-test to compare the thresholds between the 100%-validity condition and the neutral (no-expectation) condition.

### RESULTS

Figure 2 shows duration thresholds of participants in each validity condition. We conducted 2 (congruency: congruent, incongruent) x 2 (validity: 75%, 50%) repeated measures ANOVA to investigate the effect of expectation on duration thresholds. We found that the main effect of congruency is statistically significant (*F* (1,7) = 6.554, *p* = 0.034). However, the main effect of validity and interaction were not significant (*p >* 0.05). Next, we conducted post-hoc comparison tests to compare the thresholds of congruent and incongruent trials in each validity condition. We found that incongruent trials had longer duration thresholds than the congruent trials in the 75%-validity condition (*t* (7) = −3.85, *p* = 0.005). There was no difference between congruent and incongruent trials in the 50%-validity condition (*p >* 0.05). Finally, we conducted two-sample paired t-tests between (1) the 100%-validity and neutral conditions, (2) the congruent trials of 75%- and 100%-validity conditions, and (3) the congruent trials of 50%- and 100%-validity conditions. All three tests showed that the thresholds of the conditions were not statistically significantly different from each other (*p >* 0.05).

**Figure 2:**
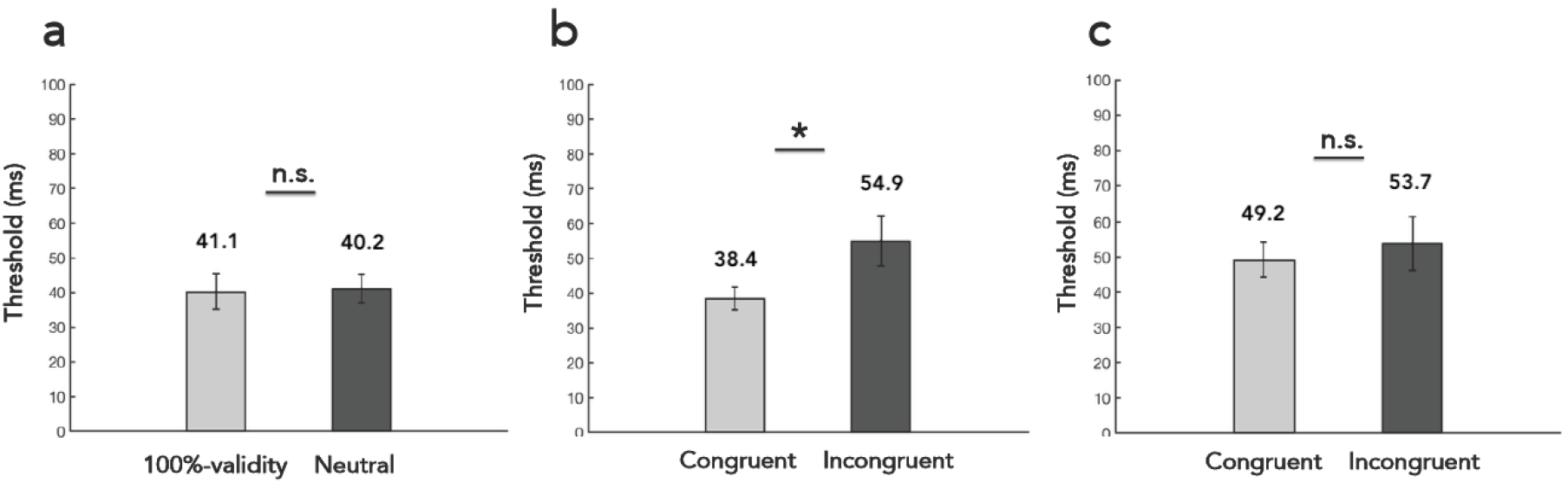
Results of the behavioral experiment. Duration thresholds of **a.** 100%-validity and neutral conditions; **b.** congruent and incongruent trials in 75%-validity condition; **c.** congruent and incongruent trials in 50%-validity condition.

### INTERMEDIATE DISCUSSION

Our behavioral results show that the thresholds of congruent (expected) and incongruent (unexpected) trials are different under the 75%-validity condition, but not under any other conditions. Specifically, unexpected stimuli led to longer duration thresholds than expected stimuli when the cue had 75% validity. This result is inline with previous findings which showed that unexpected stimulus is detected or recognized more slowly and less accurately than the expected one (Wyart et al., 2012; Stein and Peelen, 2015). Surprisingly, we also found that the thresholds of neutral-and 100%-validity conditions are not different from each other. This suggests that valid expectations do not reduce perceptual thresholds compared to the condition where there is no expectation. Also, thresholds do not differ between congruent trials of 75%- and 100%-validity conditions as well as between congruent trials of 50%- and 100%-validity conditions. Taken together, our findings suggest that *valid* expectations do *not* reduce the thresholds. Rather, the perceptual thresholds increase when the expectations are *not* met but *highly valid* for a given task.

There are two possible alternatives that may explain the underlying computational mechanism of this finding. First, it is possible that the underlying parameters of the system (*e.g.* internal noise or decision threshold) may vary based on expectation (congruency) and/or its validity. Specifically, in congruent and incongruent trials internal parameters of the system may be different so that incongruent trials are detected in longer duration than the congruent trials (De Loof et al., 2016). Alternatively, it is possible that in incongruent trials further processing may be required to make a decision, because prediction and the actual input disagree. The standard psychophysical analysis alone cannot inform as to which of these alternatives better explains the behavioral results. In order to test these alternatives we introduce a Bayesian computational model explained next.

## Modeling

Here we implement a recursive Bayesian updating scheme, in which the prior is not fixed but updated at each iteration, to model the behavioral results. By manipulating the system’s underlying parameters in different models we tested whether expectation has an effect on the underlying parameters. To test the alternative possibility, namely to test whether further processing is required in incongruent trials we compared the number of iterations calculated in congruent and incongruent trials.

### IMPLEMENTATION OF THE BAYESIAN MODEL

We used a generative model for which Bayesian inference equations were derived (see also Bitzer et al. (2014)). Figure 3 shows a schematic of the components of the Bayesian model (for a trial with 75%-validity) adapted to the present study’s experimental paradigm. As can be seen in Figure 3 the calculations were done separately for the observation on the left side and right side of the screen in each trial of the experiment.

**Figure 3:**
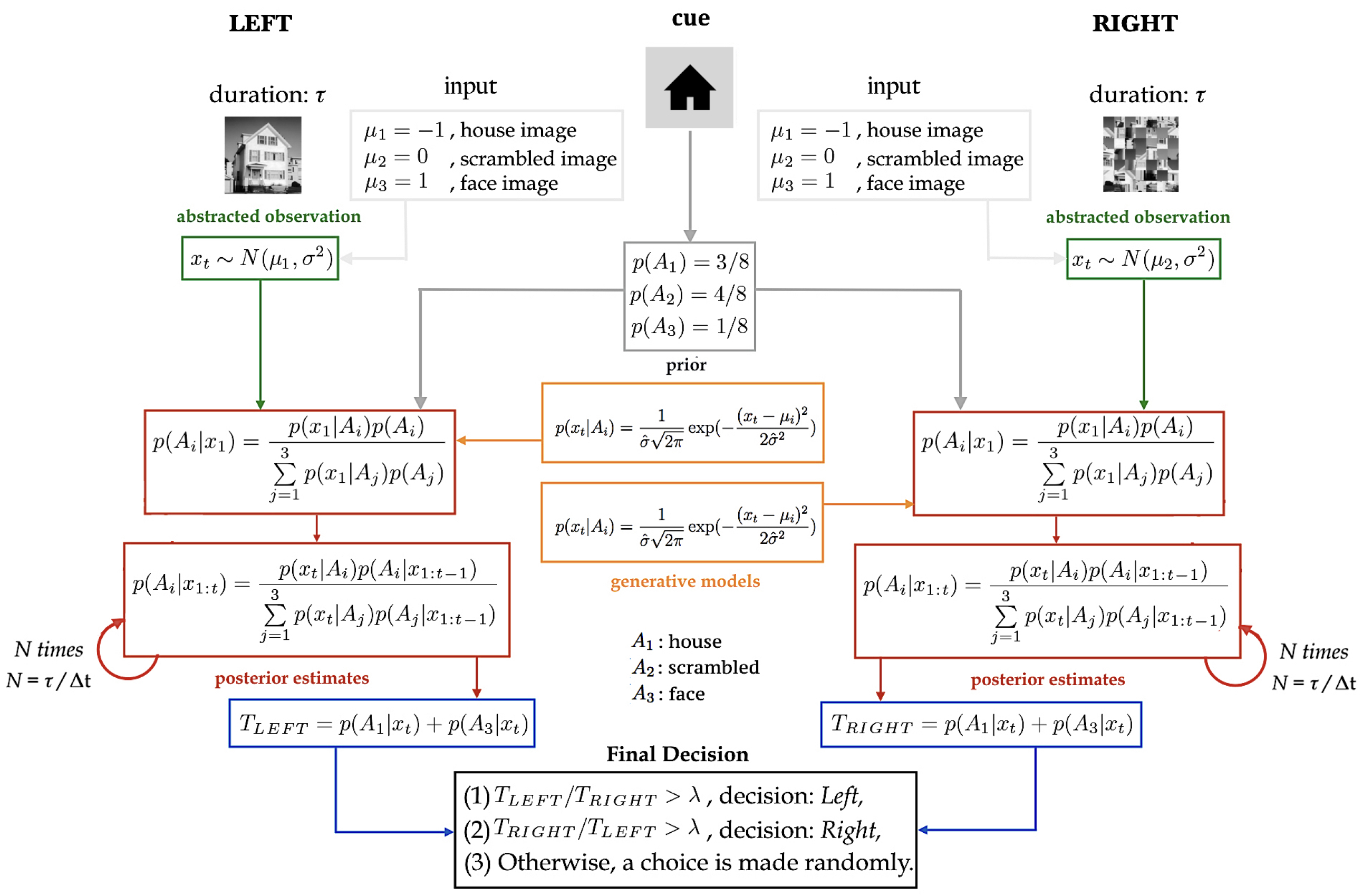
Bayesian model adapted to the current experimental paradigm. Figure shows the model for a 75%-validity condition trial as it is reflected by the values in the prior box above. See text for details.

We first defined feature values for the *input* (light gray boxes in Figure 3)

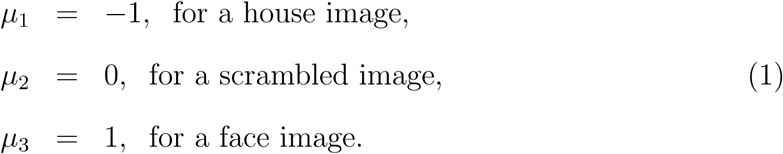

These would be the abstracted values received by the system if there were no noise. Next, it is postulated that the *abstracted observation* extracted by the system, *x*_*t*_, is drawn from a normal distribution with the corresponding *µ*_*i*_ as follows:

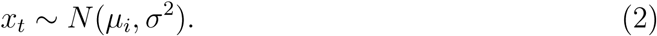

In each trial we calculated *x*_*t*_ based on the presented images on the corresponding sides. Next, we defined *generative models* for each decision alternative, *A*_*i*_: *A*_1_ for house, *A*_2_ for scrambled, and *A*_3_ for face-image. We calculated the likelihood of *x*_*t*_ under each decision alternative as

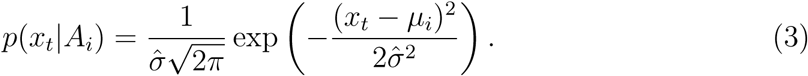

We then defined the *initial* values of the *priors* as indicated by the dark gray box in Figure 3. In each trial we defined the prior probability of observing a house-, scrambled-, and face-image:

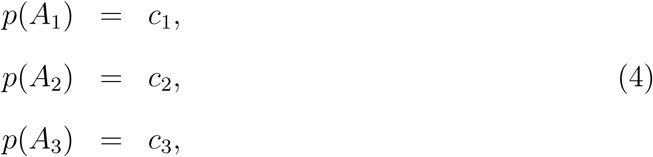

where *c*_1_, *c*_2_, and *c*_3_ are defined based on the cue validity (i.e. 100%, 75%, 50%), *and* the cue presented at each trial (i.e. face or house). For example in a trial under the 75%-validity condition if the cue is a *face* then the priors are

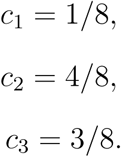

If cue is a *house*, then

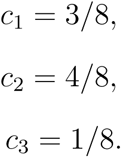

Next, we combined the likelihoods with the priors to compute *posterior estimates* for each decision alternative for both sides as follows

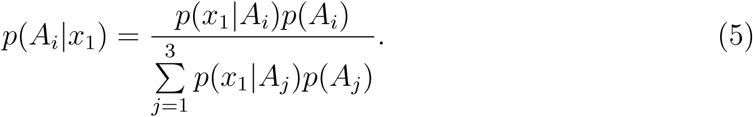

Within a single trial posterior estimates are updated recursively over time (*N* times: number of iterations) until a decision is made by the model

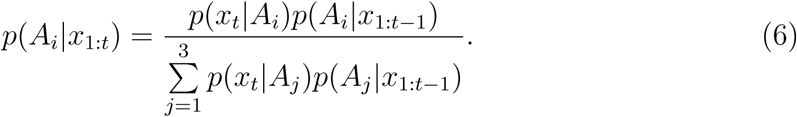

Note that, this amounts to using priors that are not fixed but updated in each iteration: posterior of the previous iteration becomes the prior for the next iteration. Number of iterations, *N*, in a single trial is determined by

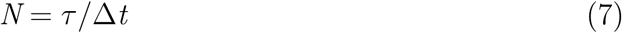

where *τ* represents the duration of presentation of the target images in this particular trial, and Δ*t* defines how long each iteration lasts in the system. Next, we calculated *probability of observing an intact image (target stimulus: face or house)* for both sides, *T*_*LEFT*_ and *T*_*RIGHT*_, by calculating the sum of last posterior of face-image and house-image as shown in blue boxes in Figure 3. At the last step, a *final decision* is made by the model using the criteria shown in black box in Figure 3. Specifically, the ratio of *T*_*LEFT*_ to *T*_*RIGHT*_ is compared to the *decision threshold*, *λ*. This evaluation determines whether the model decides *left* or *right*. If this criteria cannot be met, then a decision is made randomly.

### MODEL SIMULATIONS FOR INDIVIDUAL DATA

There were three free parameters in the model; *λ* (decision threshold), Δ*t* (how long each iteration lasts in the system), and 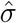 (the internal uncertainty of the decision makers representation of its observations, Eq.3). Using the optimized parameter values (that minimize the error between the model’s prediction and the real data) we ran 1000 simulations of the model to ensure the stability of the model’s predictions (Ritter et al., 2011) for each participants data. We generated separate models for 100%-, 75%-, 50%-validity conditions, and the model simulations were compared to the data of these validity conditions for each participant. Note that there was only a single difference between the models of different validity conditions, and it was the initial values of the priors (See gray box in Figure 3). Also note that there was no explicit (informative) cue in the neutral condition, which made it inherently different than other conditions. Therefore the neutral condition was not included in the simulations.

### MODEL COMPARISON

To test the first possible alternative, that is *whether underlying parameters of the system differ in different trial types*, we defined two models: in the *restricted model* a single set of parameters (3 parameters: *λ*, Δ*t*, and 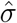) was optimized for all validity conditions and trial types (all trials in 100%-, 75%-, 50%-validity conditions) for each participant. In the *unrestricted (free) model* 5 different sets of parameters (15 parameters: 3 parameters x 5 conditions (100%, 75%-congruent, 75%-incongruent, 50%-congruent, 50%-incongruent)) were optimized; one for each trial type and each validity condition for each participant.

Next, for each participant’s data we performed a nested hypothesis test to see whether the unrestricted model models the empirical data better than the restricted model. For this aim, we performed a chi-square nested hypothesis test. Under the null hypothesis, twice the difference between the log-likelihoods of the two models has an approximate chi-square distribution, with degrees of freedom equal to 12, which is the difference in the number of parameters between the two models. Thus we reject the null hypothesis if

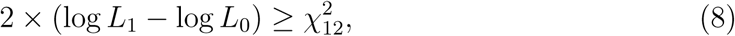

where the likelihoods *L*_0_ and *L*_1_ are calculated for the restricted and unrestricted model respectively. Note that *L* is defined as

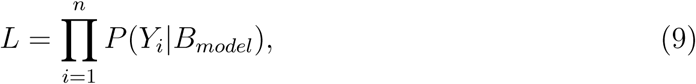

where *n* is equal to the total number of trials in each experimental condition, *Y*_*i*_ corresponds to the participant’s response in each trial, and *B*_*model*_ corresponds to the model’s prediction at each duration presentation level.

### RESULTS

Figure 4, 5, 6 and 7 show Bayesian model simulations of all validity conditions and trial types for restricted and unrestricted model for each participant. It is clear that our Bayesian scheme can successfully capture the pattern observed in the empirical data. Similar to the results of 75%-validity condition in psychophysical findings, Bayesian simulations of incongruent trials (in both models) are also shifted to the right (*i.e.* longer duration thresholds) compared to the congruent trials. This shows that the Bayesian model, just as the human participants, require a longer duration to detect the location of the intact image in an incongruent trial. Moreover, under the 50%-validity condition there is not such a clear shift, again just as in human data.

**Figure 4:**
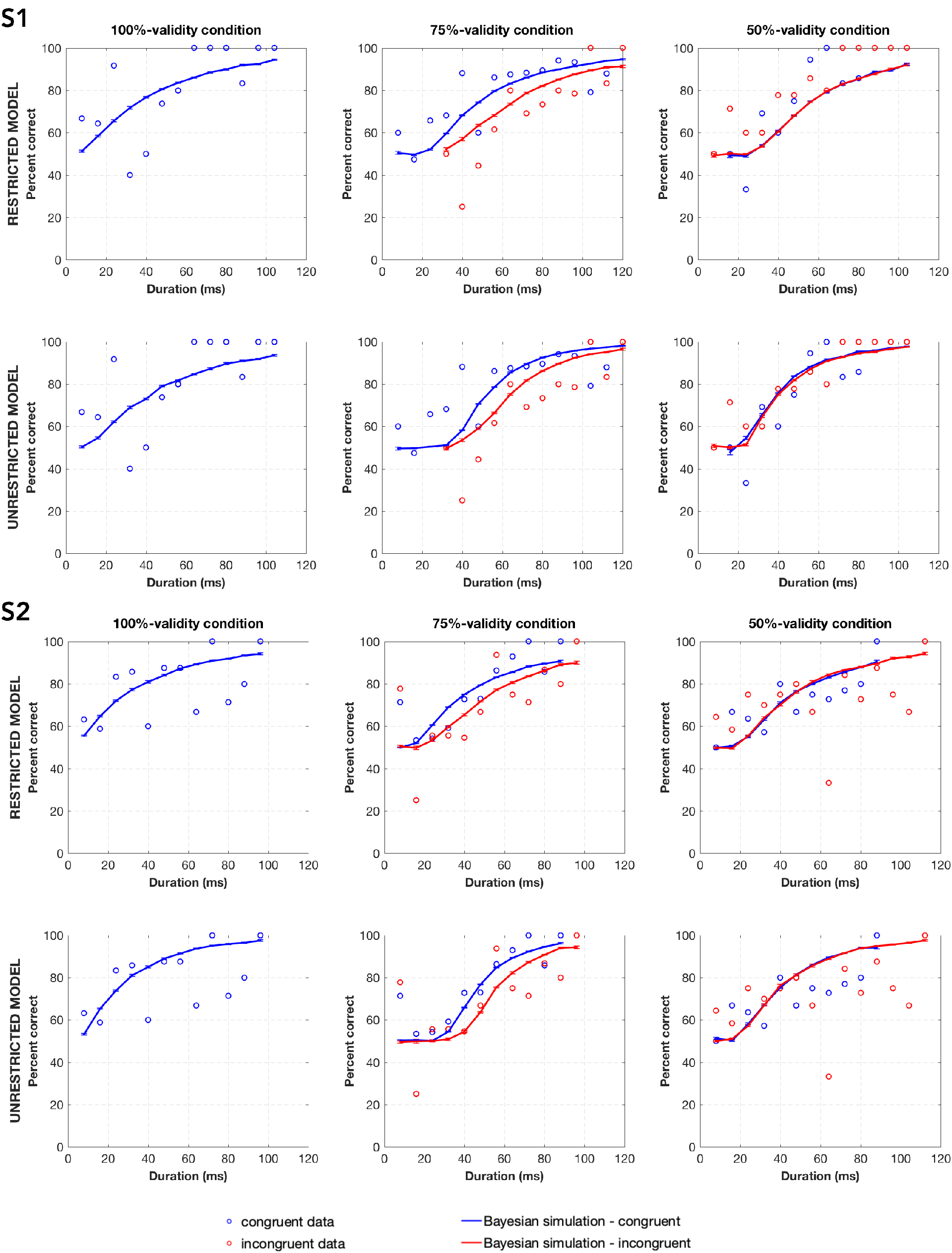
Bayesian model simulations of restricted and unrestricted model for participant 1 and 2.

**Figure 5:**
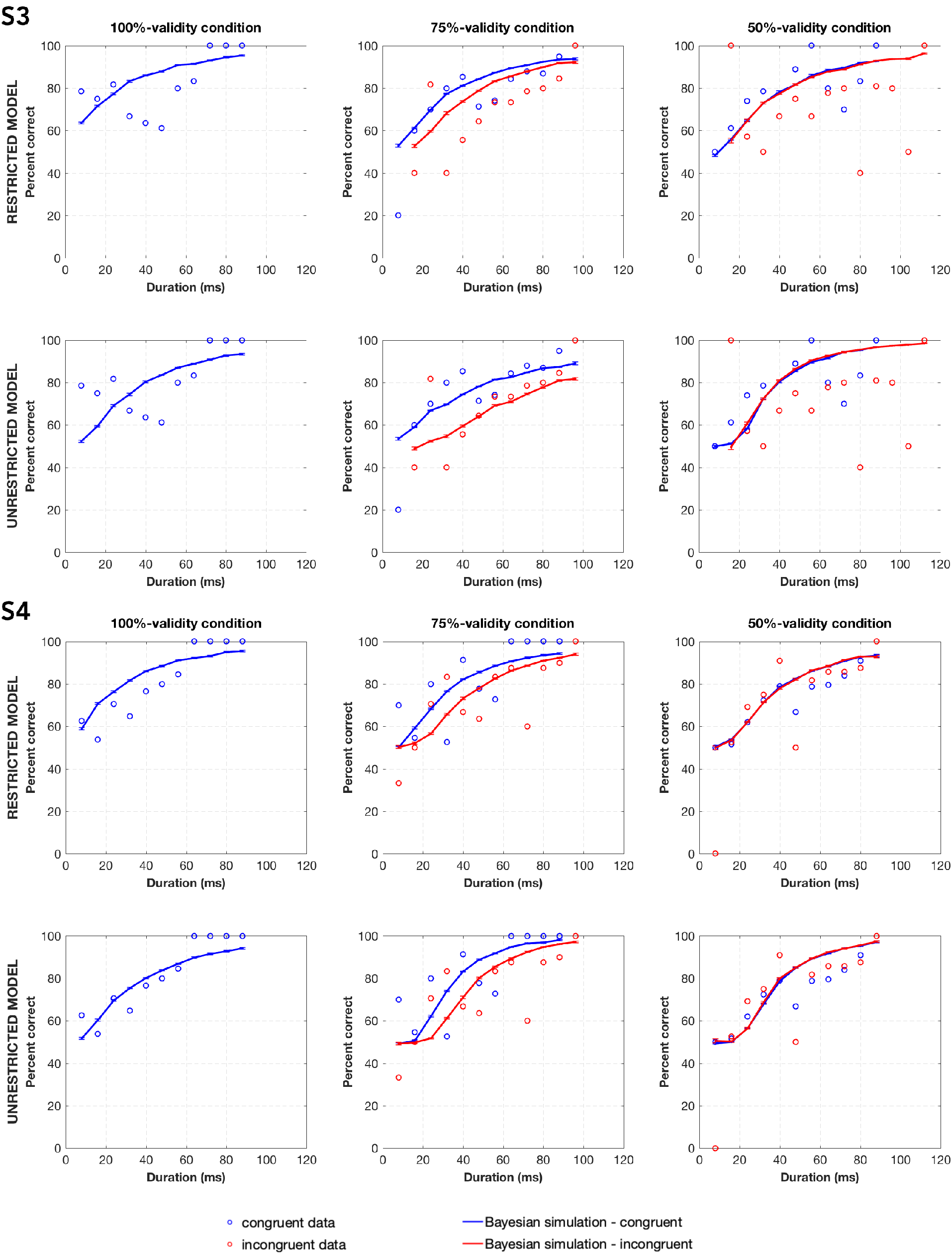
Bayesian model simulations of restricted and unrestricted model for participant 3 and 4.

**Figure 6:**
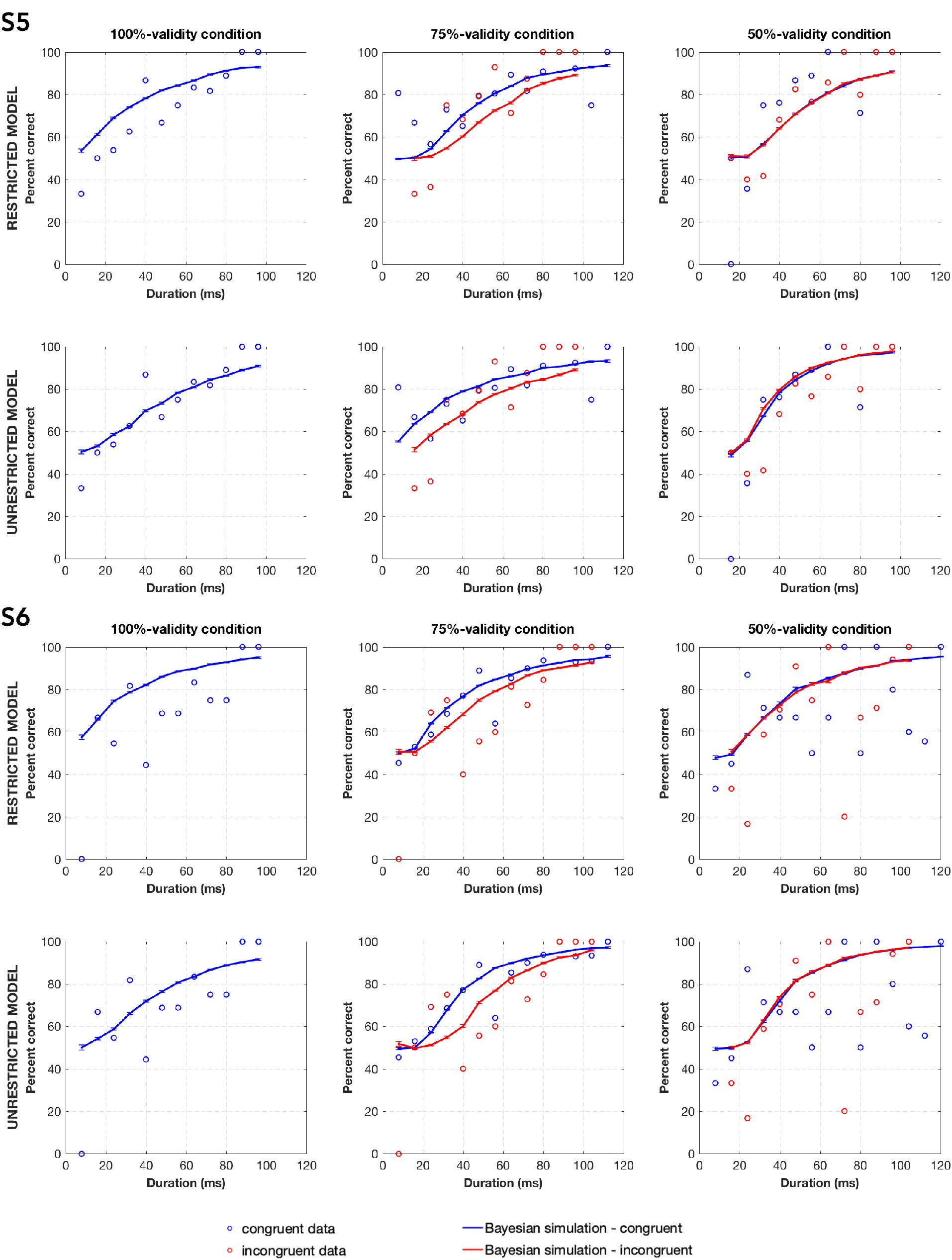
Bayesian model simulations of restricted and unrestricted model for participant 5 and 6.

**Figure 7:**
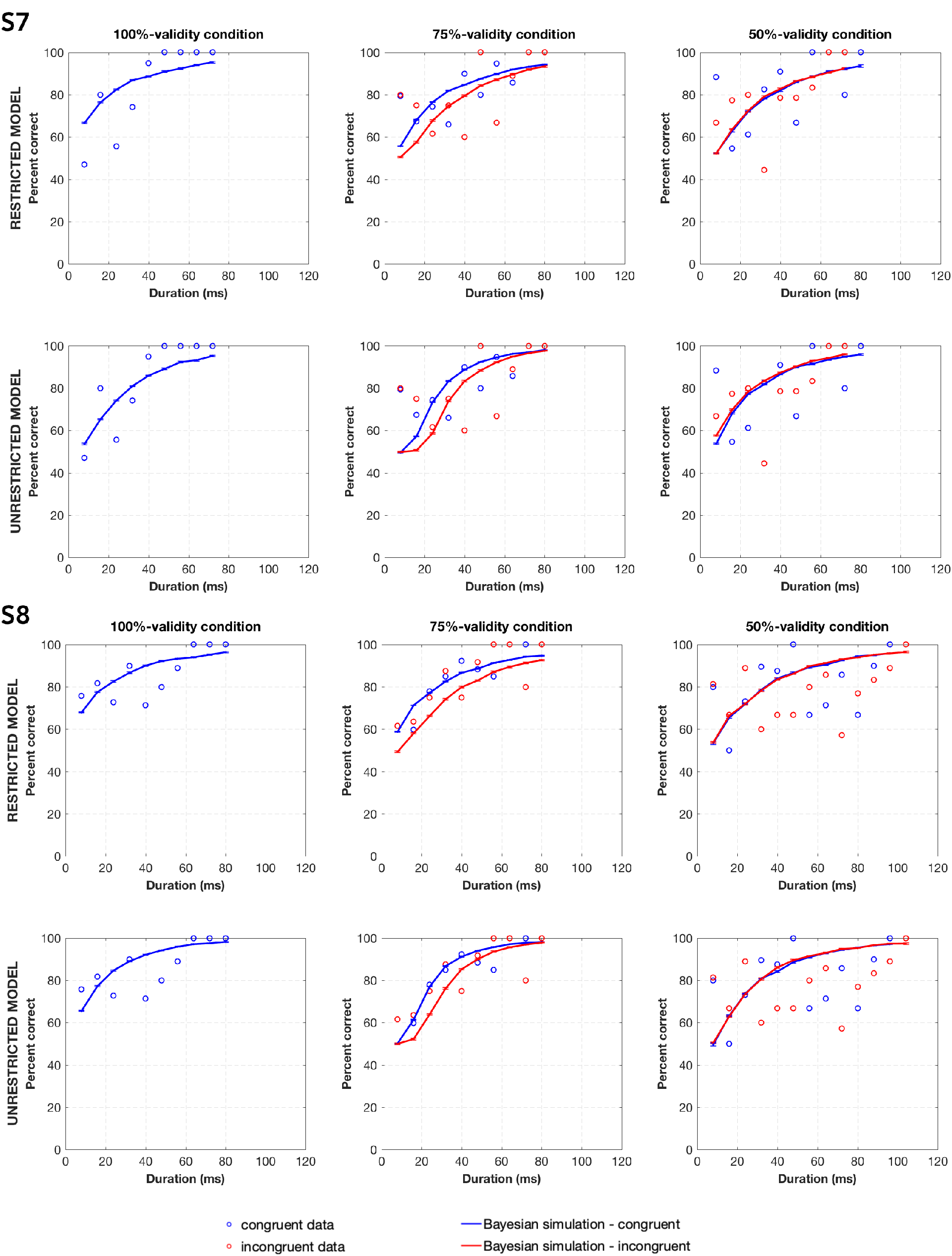
Bayesian model simulations of restricted and unrestricted model for participant 7 and 8.

The results of the likelihood-ratio tests showed that the two models are not different from each other in any participant (*p >* 0.05). This suggests that the internal parameters (λ, Δ*t*, 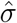) do not change with congruency (trial types: congruent, incongruent) and/or validity. This speaks against the first alternative to explain the human data we postulated earlier.

If the second alternative, that is if the system requires more time to process the visual input under the incongruent trials then the number of iterations that are needed to make a decision would be larger in those trials. To test this alternative, we calculated the number of iterations computed by the (restricted) model in congruent and incongruent trials in all validity conditions. Figure 8 shows results of number of iterations performed (posteriors computed) in each validity condition and trial type.

**Figure 8:**
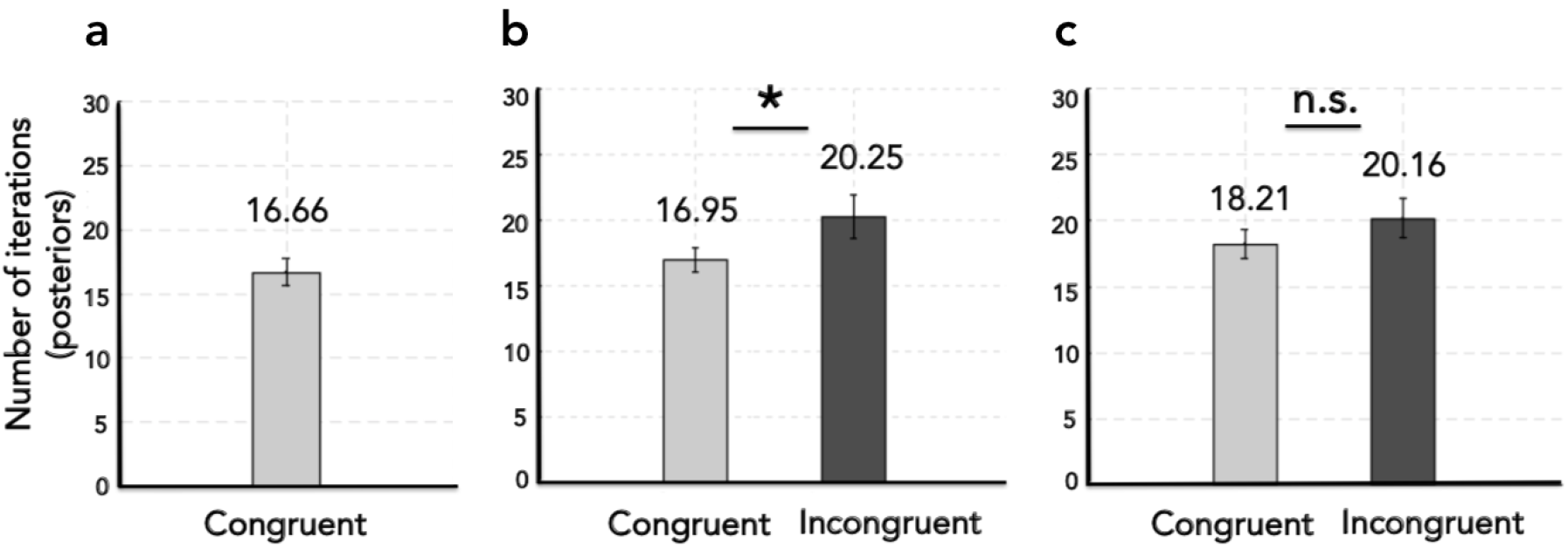
Number of iterations, N (posterior computations) in congruent and incongruent trials in all validity conditions. a. 100%-validity condition. b. 75%-validity condition. c. 50%-validity condition.

We performed a 2 (congruency: congruent, incongruent) x 2 (validity: 75%, 50%) repeated measures ANOVA to investigate the effect of congruency and validity on number of iterations (posteriors). As expected, the main effect of congruency was significant (*F* (1,7) = 11.731, *p* = 0.011). However, the main effect of validity and interaction were not significant (*p >* 0.05).

Next, we performed post-hoc comparison tests to see whether the number of iterations differ based on congruency speficially in each validity condition. In 75%-validity condition number of posteriors computed in incongruent trials are higher than the congruent trials (*t* (7) = −3.4798, *p* = 0.0103). However, there was no difference between congruent and incongruent trials in 50%-validity condition (*p >* 0.05). Also, there was no difference in number of iterations between (1) congruent trials of 50%- and 100%-validity conditions as well as between (2) congruent trials of 75%- and 100%-validity conditions (*p >* 0.05). Overall these results agree with the behavioral data remarkably well.

## General Discussion

In this study we investigated the effect of expectation on early visual processing by measuring perceptual thresholds. To this aim, we systematically manipulated the expectation validity in different experimental conditions, and measured duration thresholds to examine whether the perceptual thresholds to detect a stimulus vary depending on expectation and/or its validity. We then presented a recursive Bayesian updating scheme to elucidate the underlying mechanisms of the findings we observe in the behavioral experiment. Previous findings already showed that under several circumstances human behavior is nearly Bayes-optimal. And it is also clear that perceptual decision-making processes are strongly influenced by expectations. However, our study goes beyond these findings because, to our knowledge, this is the first study that systematically investigates the behavioral effect of expectation on *early visual processes* at the threshold level, and unravels possible computational mechanisms underlying the behavioral results using a *recursive* Bayesian updating scheme.

### Expectation affects visual perceptual thresholds only when the expectation validity is relatively high

Our behavioral results showed that unmet expectations can shape early visual processes only when the cue has a relatively high validity (*i.e.* 75%). In a similar individuation task, under 80%-validity and neutral conditions, De Loof et al. (2016) showed that expectations speed up perceptual *decisions* by measuring response times (RT), which reflect the time required by a combination of early visual, cognitive and decision-making processes to give a response. Our study furthers these findings and shows that not only the perceptual decisions, but even the early visual processes in isolation are affected by expectations. Furthermore, surprisingly, we found no difference in perceptual thresholds of 100%-validity and neutral conditions as well as congruent trials of 75%- and 100%-validity condition, and congruent trials of 50%- and 100%-validity condition. Taken together, our findings show that the perceptual thresholds do not decrease if a stimulus is expected, rather that the thresholds increase if expectations are not met, specifically when those expectations were high.

### Unexpected stimulus leads to further processing, rather than a change in the internal parameters of the system

The behavioral findings above led us to consider two non-mutually exclusive possibilities that may explain the observed results. First, internal parameters of the system (*e.g.* the decision threshold) may differ with expectation and/or its validity. Second, further processing may be required to make a decision when the expectations are not met. To test these hypotheses, we used a recursive Bayesian updating scheme where the prior is not fixed but updated at each iteration to model our behavioral findings under the 100%-, 75%-, and 50%-validity conditions.

First, to examine the first alternative, we performed Bayesian simulations using a restricted model and a unrestricted model, and compared how successfully they fit the empirical data. In general, our findings on both models revealed that our recursive Bayesian scheme can capture the pattern observed in the empirical data. Specifically, as in the psychophysical results Bayesian model simulations for 75%-validity condition showed that incongruent trials are detected in longer duration than the congruent trials. However, this pattern is not observed in 50%-validity condition. This finding further supports the idea that humans behave in a Bayes-optimal fashion in which perceptual decisions are made by combining the sensory input with the prior in a probabilistic manner (Ernst and Banks, 2002; Kersten et al., 2004). Critically our model comparison analysis showed that the two models, restricted and unrestricted, are not different from each other in any participant. This shows that expectation and its validity do not modulate the underlying parameters of the system (i.e. decision threshold, internal uncertainty, how long each iteration lasts in the system), argues against the first alternative mechanism to explain the behavioral data.

Next, for the second alternative, we calculated the number of iterations computed in congruent (expected) and incongruent (unexpected) trials. Our results showed that more number of iterations are calculated in incongruent trials only when the expectation’s validity is relatively high (75%). This reveals that in order to make a decision more posteriors should be updated in incongruent trials, which is an indicator of further processing within a single trial. This finding is remarkably consistent with our behavioral findings and suggests that the observed increase in the perceptual thresholds of incongruent trials in 75%-validity condition is due to an additional processing rather than a change in the system’s internal parameters.

In a study that we introduced earlier, De Loof et al. (2016) studied the response times using a similar experimental design. In that study, the authors used drift-diffusion model (DDM) (Ratcliff, 1978) to model the empirical results, and found that the unexpected stimuli led to increased boundary separation parameters, which is defined as the internal threshold that is required to reach a decision (De Loof et al., 2016). This result appears at odds with our findings. We argue that, even though the DDM model is a well-studied and highly useful model to understand the underlying processes in perceptual decisions, it does not capture certain characteristics of the current experimental paradigm because, for example, the validity of the expectation or more importantly the temporal dynamics throughout a trial cannot be modeled with DDM (Huk et al., 2018). On the other hand, our Bayesian scheme provides us the opportunity to (1) define task-irrelevant prior, (2) set its validity, and (3) recursively update posterior estimates within a single trial considering that perception is a *dynamic* inference process. Indeed, when we performed a DDM analysis using the behavioral threshold values, we found that the model estimates of the boundary separation increased under the incongruent conditions (analyses and results not reported here). This outcome further shows that the recursive Bayesian model captures some important details about the dynamics of the underlying processes that the DDM model cannot.

### Cortical Models, Predictive Coding: Do expectation violations require prediction error computation of the ‘error units’?

Bayesian approaches to understand the brain function, not only behavior, have gathered considerable support in the literature. Accordingly, several generic models for brain function have been proposed, which have computational concepts that are analogous to the ones in Bayesian framework (Rao and Ballard, 1999; Friston, 2005; Heeger, 2017). Several fMRI studies showed that there is an increased BOLD activity in response to an unexpected stimulus compared to an expected one (Yoshiura et al., 1999; Marois et al., 2000; Kok et al., 2011). One interpretation of this finding is that it reflects the *prediction error* signal of the *error units* (Summerfield and Egner, 2009; *Kok et al., 2011), which are introduced in the predictive coding theory (Friston, 2005).* Predictive coding theory (PCT) proposes a generic model for brain function and posits that based on prior information the brain computes internal predictions about the upcoming sensory input. When there is a mismatch between the predictions and the sensory input, a mismatch signal is computed which is called the prediction error. The PCT postulates that the prediction- and the error signals are computed by specific neuron units called representation units (RU) and error units (EU) respectively, which are hypothesized to exist at each level of the cortical hierarchy. In the case of a mismatch, the error signal is conveyed to the higher levels of the cortical hierarchy to update the predictions. If there is a match between the predictions and the sensory input, then the error neurons do not respond vigorously, which is interpreted as the silencing of prediction error. Therefore, the PCT posits that the information processing in the brain is a dynamic interplay between the prediction signals by RU and prediction error signals by EU, which are conveyed by feedforward and feedback connections. Even though a growing number of studies suggest that the observed increase of BOLD response is an indicator of the prediction error signal of the error units, there is still no empirical evidence that reveals the existence of the error units in the cortex.

Alternative to PCT, the cortical function theory of Heeger (2017) posits that the whole process can be executed without the existence of specific error- and representation units posited by the PCT. Heeger (2017) claims that the information processing can be accomplished and explained only by feedforward-feedback connections in the brain. The findings of the current study show that in the case of a mismatch condition further processing is required to make a decision. We suggest that further processing may be an indicator of a change in the feedforward-feedback connections during information processing. When an unexpected stimulus is presented, an additional processing may be required compared to an expected stimulus presentation, and this processing can be implemented with additional feedforward and feedback interactions. In this sense the brain does not need to have separate “error units” that compute prediction error as posited by predictive coding models (Friston, 2005). Rather, the same processing may be implemented with additional computations to process an unexpected stimulus compared to the expected one.

In short, we suggest that an increase in BOLD response to an unexpected stimulus does not necessarily reflect the “error unit” activity posited by the PCT. Rather, it may indicate an additional processing of the neural populations via feedforward-feedback connections as suggested by Heeger (2017), and this would be consistent with our findings on human behavior and Bayesian model. It should be noted that our Bayesian scheme is useful to investigate why there is such a difference between the perception of expected and unexpected stimuli. However, it does not reveal how this process can be executed at the level of cortex. Predictive computation models (Friston, 2005; Heeger, 2017) should be employed to empirically test how this is accomplished, and it is the subject for future research.

## Conclusion

In summary, our results offer several surprising and interesting results about the role of expectation on early visual processing. Firstly, we showed that expectations do not make the participants faster, rather unmet expectations make them slower. Secondly, using a simple and parsimonious model we found that this slow-down in human behavior can be explained by further processing required in the visual system when the expectations are violated. Furthermore, the experimental paradigm and the computational model introduced here have the potential to be expanded and used for new and novel studies.

## Acknowledgements

This work was funded by a grant of the Turkish National Scientific and Technological Council (TUBITAK 217K163) awarded to HB.

## Footnotes

1 To control for any possible confounding effects of training, we conducted a control experiment on a separate group of participants who did not participate in a training session prior to the experimental session. See Supplementary Material for methodological details of the experiment. Also see Supplementary Figure S1 for results of the control experiment.

