## Supplementary Material for "When expectations are not met: unraveling the computational mechanisms underlying the effect of expectation on perceptual thresholds"

---

### Control Experiment

To eliminate any confounding effect of training on our behavioral findings, we conducted the same experiment on a separate group of participants who did not participate in a training session before the main experiment.

### Methods

*Participants*

10 participants (Group 1: 6 participants, $26 \pm 1.78$ years; Group 2: 4 participants, $26.25 \pm 2.22$ years ) participated in the control experiment. All participants had normal or corrected to normal vision and reported no history of neurological disorder. Informed consent was taken prior to the experiment. They were randomly assigned to one of the two experimental groups (See *Procedure* below for details). The experiment was approved by the Research Ethics Committee at Bilkent University.

*Stimuli, Experimental Design and Procedure*

All stimuli and experimental design were exactly the same as in the main experiment. Participants were randomly assigned to one of the two experimental groups: Group 1 completed two experimental conditions that comprised 75%-validity and neutral conditions, and Group 2 completed 50%-validity and neutral conditions. All participants completed each experimental condition in random order in separate sessions.

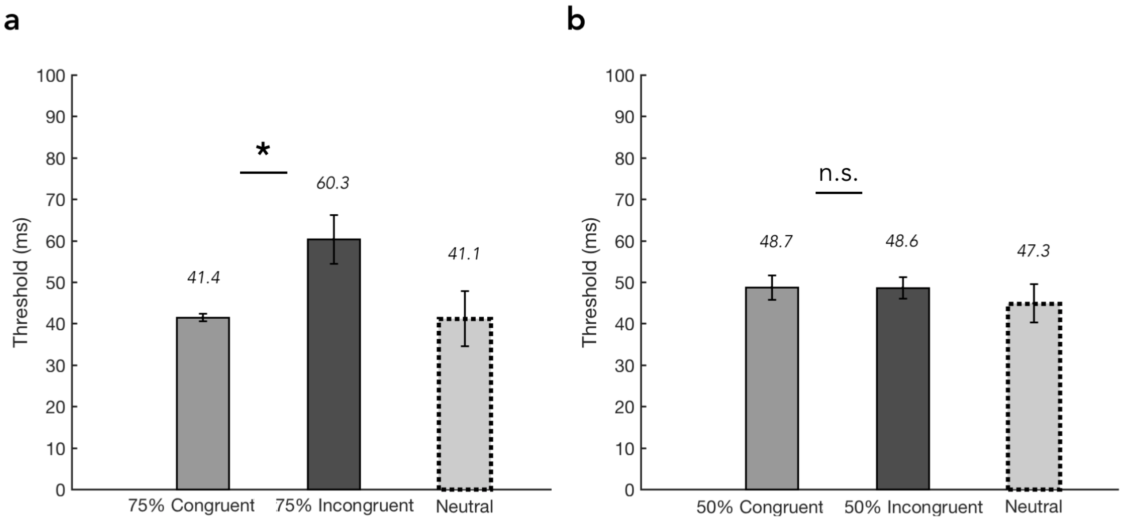


**Figure S1. Results of the control experiment.** a. Results of Group 1 who completed 75%-validity and neutral conditions. b. Results of Group 2 who completed 50%-validity and neutral conditions.

### Results

We conducted two-sample paired t-tests separately for Group 1 and Group 2 to see whether duration thresholds of congruent and incongruent trials differ. Results of the control experiment can be seen in Figure S1. Results of Group 1 showed that in 75%-validity condition there was statistically significant difference between congruent and incongruent trials in duration thresholds ($t(5)$=-2.8833, $p$<0.0345). Results of Group 2 showed that in 50%-validity condition there was not a significant difference between congruent and incongruent trials in duration thresholds ($p > 0.05$). Also, thresholds of neutral condition were not different from congruent trials of 75%- (Group 1) and 50%- validity (Group 2) conditions ($p > 0.05$). Taken together, results of the control experiment confirmed that the findings of the main experiment are not due to an effect of training.
